# Gut Dysbiosis with a Pathobiont Shifts the Intestinal Microbiota Profile and Accelerates Lupus Nephritis

**DOI:** 10.1101/2020.10.12.310763

**Authors:** Giancarlo R. Valiente, Armin Munir, Marcia L. Hart, Perry Blough, Takuma T. Wada, Emma E. Dalan, William L. Willis, Lai-Chu Wu, Aharon G. Freud, Wael N. Jarjour

**Author notes:** **CORRESPONDING AUTHOR**: Wael N. Jarjour.

## Abstract

The gut microbiota (GM) exerts a strong influence over the host immune system and dysbiosis of this microbial community can affect the clinical phenotype in chronic inflammatory conditions. To explore the role of the GM in lupus nephritis, we colonized NZM2410 mice with Segmented Filamentous Bacteria (SFB). Gut colonization with SFB was associated with worsening glomerulonephritis, glomerular and tubular immune complex deposition and interstitial inflammation compared to NZM2410 mice free of SFB. With SFB colonization mice experienced an increase in small intestinal lamina propria Th17 cells and group 3 innate lymphoid cells (ILC3s). However, although serum IL-17A expression was elevated in these mice, Th17 cells and ILC3s were not detected in the inflammatory infiltrate in the kidney. In contrast, serum and kidney tissue expression of the macrophage chemoattractants MCP-1 and CXCL1 were significantly elevated in SFB colonized mice. Furthermore, kidney infiltrating F4/80+CD206+ M2-like macrophages were significantly increased in these mice. Evidence of increased gut permeability or “leakiness” was detected in SFB colonized mice. Finally, the intestinal microbiome of SFB colonized mice at 15 and 30 weeks of age exhibited dysbiosis when compared to uncolonized mice at the same time points. Both microbial relative abundance as well as biodiversity of colonized mice was found to be altered. Collectively, SFB gut colonization in the NZM2410 mouse exacerbates kidney disease, promotes kidney M2-like macrophage infiltration and overall intestinal microbiota dysbiosis.

## INTRODUCTION

Our understanding of the intestinal microbiota’s role in autoimmune disease is rapidly evolving. There is a paucity of knowledge concerning the specific impact of the GM in Systemic Lupus Erythematosus (SLE) and more specifically lupus nephritis (LN). Studies have observed GM dysbiosis in patients with lupus but its importance in disease pathogenesis and exacerbation are not fully appreciated.

The intestinal microbiota plays an important role in many aspects of normal homeostasis and disease states. Early studies focused on the role of the GM in gastrointestinal disease (1–3). Evidence of the extraintestinal effects that enteric bacteria can exert on host has been documented in the past (4). One of the most striking examples is that of *Tropheryma whipplei*, the bacterium that causes Whipples’ disease (5). Patients infected with *T. whipplei* can develop widespread arthritis, chronic diarrhea and even central nervous system and cardiopulmonary manifestations (6). Attention grew when observations of gut dysbiosis were made in non-intestinal autoimmune diseases such as type I diabetes (7). More recently, the GM has been implicated in the pathogenesis of inflammatory arthritis in rodents and humans (8–10). With respect to lupus and LN, only a small number of studies have been performed in humans or mice (11–16). Identification of specific commensal organisms that become pathogenic in a disease context, also known as “pathobionts”, have not yet been studied in LN. Pathobionts in autoimmune diseases like inflammatory arthritis have been shown to exhibit immune-activating properties (9, 17).

Insight into this complex relationship has been studied in mice by utilizing the commensal organism Segmented Filamentous Bacteria (SFB). Germ-free (GF) wildtype mice do not possess fully formed gut-associated lymphoid tissues (GALT) and have deficiencies in the myeloid lineage (18–21). The downstream consequences on the immune system include a significant reduction in the intestinal Th17 repertoire as well as having nearly undetectable IL-17A and IL-22 (22). Monocolonization of germ-free wildtype mice with SFB induces Th17 cell differentiation and proliferation that resembled specific-pathogen free (SPF) raised wildtype mice (22, 23). Although no pathogenic effect was observed in wildtype mice following SFB colonization, most of the expanded Th17 cells were SFB-specific and required CD11c^+^ myeloid cells for proper SFB-antigen presentation (24). Furthermore, macrophages were found to be essential in the process of SFB-induced Th17 cell differentiation and expansion (25). Interestingly, studies have shown that SFB is a potent pathobiont in mouse models of inflammatory arthritis wherein intensified erosions and elevated autoantibody titers are seen in SFB colonized mice (17, 26–28). However, distinct GM species that directly affect lupus or LN have yet to be identified in animal models or patients with lupus.

We chose to study the effects of the commensal pathobiont, SFB, in the NZM2410 lupus nephritis mouse model due to robust development of glomerulonephritis and eventual mortality from immune-mediated renal dysfunction, a prominent feature in up 25-60% of patients with SLE (29). In this study we report that following colonization with SFB, mice developed severe membranoproliferative glomerulonephritis marked by immune complex deposition and M2-like macrophage cell infiltration. Small intestinal Th17 cells and ILC3s were also expanded in these mice when compared to controls. Evidence of increased intestinal permeability in SFB inoculated mice was detected in the small intestine. Microbiome 16S ribosomal RNA (rRNA) analysis revealed GM dysbiosis in SFB inoculated mice. Furthermore, altered gut microbiome composition was evident in both early and late disease and when compared to control mice. Interestingly, several phylogenetic changes in the microbiota were detected in SFB-colonized mice displaying changes that have previously been observed in both patients and mouse models. These findings suggest that gut dysbiosis introduced by intestinal pathobionts such as SFB can influence inflammation at distant tissue sites in autoimmune conditions like LN.

## Materials and Methods

### Ethics Statement

This study was carried out in strict accordance with the recommendations in the Guide for the Care and Use of Laboratory Animals of the National Institutes of Health. The protocol was approved by the Institutional Animal Care and Use Committee (IACUC) of The Ohio State University (Animal Welfare Assurance Number: A3261-01). All animal experiments were conducted under IACUC protocol #2017A00000032. For anesthesia and euthanasia, isoflurane and CO2 were used, respectively, according to the IACUC protocol.

### Mice

NZM2410/J (NZM2410) and +/- SFB-colonized C57BL/6 mice were purchased from The Jackson Laboratory (Bar Harbor, ME) and Taconic Farms (Rensselaer, NY) respectively. Upon arrival at our vivarium, all mice were tested for SFB gut colonization using an SFB-specific primer during polymerase chain reaction testing (PCR). Mice were housed in a specific pathogen free (SPF) facility and maintained under protocols approved by the Institutional Animal Care and Use Committee (IACUC) at The Ohio State University Wexner Medical Center (OSUWMC). The vivarium facility was sustained at 22-23 °C and 30-50% relative humidity with a 12 hour light/dark cycle. Chow and water were supplied ad libitum. Feces from +SFB and –SFB colonized Taconic C57BL/6 mice were homogenized with PBS into a slurry. Oral gavage of NZM2410 mice with 150 μL of fecal slurry was performed to inoculate mice with SFB (n = 8) or without SFB (n = 8). Serum was analyzed for blood urea nitrogen (BUN) levels in all NZM2410 mice at 30 weeks of age. At the same time tissues were harvested for downstream analyses in which 3-4 +SFB mice and 3-4 -SFB mice were randomly chosen.

### BUN measurements

Whole blood from NZM2410/J mice was obtained via submandibular bleeds. Serum was then isolated by spinning whole blood in Microtainer Serum Separator Tubes (BD,). MaxDiscovery Blood Urea Nitrogen Enzymatic Assay Kits were used to quantify BUN via colorimetric assay (Bioo Scientific Corporation, Austin, TX) according to the manufacturer’s protocol. Blood Urea Nitrogen (BUN) levels were measured at 30 weeks of age; BUN levels above 50 mg/dL were defined as a clinical indication of progressive kidney damage.

### SFB Polymerase Chain Reaction (PCR)

Fecal pellets from each mouse were initially stored at −20 °C. Fecal DNA was isolated using a DNA Stool Mini Kit (Qiagen, 51504). To detect presence or absence of SFB, the SFB-specific primers GACGCTGAGGCATGAGAGCAT and GACGGCACGGATTGTTATTCA were used (30). To resolve the DNA bands, ethidium bromide and 2% agarose gels were utilized and visualized using a UV lamp.

### Fecal DNA extraction and purification

Fecal pellets were placed in a sterile round-bottom tube and stored at −20 °C until processed for DNA extraction using the QIAamp DNA Stool Mini Kit (Qiagen, Germantown, MD). DNA concentrations determined spectrophotometrically (Molecular Devices, San Jose, CA) and fluorometrically (Qubit dsDNA BR assay, Life Technologies, Carlsbad CA). Purified DNA samples were stored at −20 °C until 16S rRNA sequencing.

### Histopathology

Formalin fixed paraffin-embedded (FFPE) tissues were sectioned at 4 μm, placed on slides and stained with H&E, PAS or Jones’ methenamine at The Ohio State Wexner Medical Center Pathology Core Facility. Glomeruli sizes are representative of 60 glomeruli counted for each of 4 +SFB and 4 −SFB control mice. Slides were scanned with an Aperio Scanscope XT from 2X to 40X. Histopathological assessment of kidney tissue was performed by a board-certified veterinary pathologist; kidney pathology was defined as cellular proliferation, hyaline deposits, cellular crescents and protein casts. Glomerular lesions were graded on a scale of 0–3 for increased cellularity, increased mesangial matrix, necrosis, percentage of sclerotic glomeruli, and presence of crescents [27]. Similarly, tubulointerstitial lesions were graded on a scale of 0–3 for interstitial mononuclear infiltration, tubular damage, interstitial fibrosis, and vasculitis.

### ELISA

Serum was tested for antibodies, cytokines and chemokines. Conventional sandwich ELISAs were used to test for anti-dsDNA Ab (Alpha Diagnostic International, San Antonio, TX), MCP-1 and CXCL1 (Invitrogen, Carlsbad, CA). Electrochemiluminescence ELISAs (Meso Scale Diagnostics, Rockville, MD) were used to test for IL-1β, IL-6, IL-17A, TNF-α. All other autoantibodies were tested for using an autoantigen microarray panel (University of Texas, Southwestern, TX).

### Lamina propria lymphocyte isolation

Mice were euthanized and small intestine removed and placed in ice-cold PBS. After removal of residual mesenteric fat tissue, Peyer’s patches were carefully excised, and the intestine was opened longitudinally. The intestine was then thoroughly washed in ice-cold PBS and cut into 1.5 cm pieces. The pieces were incubated four times in 5 mL of 5 mM EDTA in HBSS for 15 min at 37°C with slow rotation (100 rpm) in Teflon-coated flasks. After each incubation, the epithelial cell layer, containing the intraepithelial lymphocytes (IELs), was removed by intensive vortexing and passing through a steel mesh strainer and new EDTA solution was added. After the fourth EDTA incubation the pieces were cut into 2 mm2 pieces and placed in 5 mL digestion solution containing 5% fetal bovine serum (Gibco), 10 mM HEPES (Gibco), 1 mg/mL Collagenase Type VIII (Sigma), 40 μg/mL DNase I (Sigma), and 1 mg/mL Dispase II (Sigma). Digestion was performed by incubating the pieces at 37°C for 15 min with slow rotation. After the initial 15 min, the solution was vortexed intensely and passed through a 100 μm cell strainer. Flow through was washed in HBSS supplemented with 5% FBS then and resuspended in 10 mL of the HBSS/5% FBS. The cell suspension was overlaid on top of a 30%/100% Percoll gradient in a 50 mL Falcon tube. Percoll gradient separation was performed by centrifugation for 30 min at 670g at room temperature. Lamina propria lymphocytes (LPLs) were collected at the interphase of the Percoll gradient, washed once, and resuspended in FACS buffer. The cells were used immediately for experiments.

### Surface and nuclear staining for flow cytometry

Surface staining was performed for 30 min with a corresponding cocktail of fluorescently labeled antibodies. After surface staining, the cells were resuspended in Fixation/Permeabilization solution (Foxp3/Transcription Factor Staining Buffer Set, Invitrogen), washed and then resuspended in Permeabilization Buffer and nuclear transcription factor staining was performed as per the manufacturer’s protocol.

### Flow cytometry and antibodies

Flow cytometric analysis was performed on LSR II (BD Biosciences) instrument and analyzed using FlowJo software (Tree Star Inc.). All antibodies were purchased from Miltenyi Biotec, Invitrogen or BD Biosciences.

### Fluorescent immunohistochemistry

Immunohistochemistry (IHC) was performed on mouse tissue that was formalin fixed and paraffin embedded (FFPE). Specimen slides were deparaffinized, rehydrated and then antigen retrieval was performed. The slides were incubated with primary antibodies overnight at 4°C. The primary antibodies used were rabbit anti-mouse CD 127(blah), goat anti-mouse IL-22 (blah), and a rat anti-mouse lineage cocktail containing CD3, CD4, CD11b, CD19 and F4/80. After the slides were washed, donkey anti-goat AF 555 (blah), goat anti-rabbit AF 647, and goat anti-rat AF 488 were applied. The samples were incubated with DAPI. For C3 staining, the tissues were incubated with rabbit anti-mouse C3 overnight at 4°C. Slides were washed and goat anti-rabbit IgG AF 555 () was applied. Mounting was performed using Prolong Antifade something Diamond. Images were captured using EVOS FL Cell Imaging System (Thermo-Fisher Scientific) and confocal microscopy.

### Chromogenic immunohistochemistry

Three modified IHC methods were utilized: *Proteolytic IHC modified* (FFPE), *traditional IHC modified* (FFPE) and *fresh frozen IHC modified* (fresh frozen tissue). *Proteolytic immunofluorescence modified:* FFPE kidney tissue slides were heated in an oven at 60 °C to melt the paraffin wax. The remaining paraffin was dissolved and tissues dehydrated in xylene (Millipore Sigma, Burlington, MA). Slides were then rehydrated by serial ethanol washes. Proteolytic Ag retrieval was performed via 75 μg/mL pronase (Sigma-Aldrich) in tris-buffered saline, pH 7.4 (Bio-Rad, Hercules, CA). Slides were washed and incubated with appropriate antibody and subsequently sealed with Prolong Diamond Antifade mountant (Invitrogen, Carlsbad, CA) and a #1.5 coverslip (Fisher Scientific). *Traditional IHC modified:* FFPE kidney tissue slides were initially prepared as above. Samples were dehydrated and rehydrated and then heat induced Ag retrieval. Slides were washed and incubated with appropriate antibody and subsequently sealed as above. *Fresh frozen IHC modified:* kidney and SI tissue slides were briefly dried and fixed in periodate-lysine-paraformaldehyde (PLP) at 4 °C. PLP was made in our laboratory by mixing 0.2 grams of sodium periodate (Sigma-Aldrich), 0.375M L-lysine-HCl (Sigma-Aldrich), 4% formaldehyde (Sigma-Aldrich) and 0.2M NaPO4 buffer (Na2HPO4 and NaH2PO4 salts from Sigma-Aldrich). Samples were dehydrated and rehydrated and then heat induced Ag retrieval. Slides were washed and incubated with appropriate antibody and subsequently sealed as above.

### Confocal microscopy

All confocal imaging was performed at the OSU Campus Microscopy and Imaging Facility on the Olympus FV1000 Spectral scanning laser confocal system using a 40X oil objective (UPLFLN, numerical aperture 1.3) for kidney and SI sections. Confocal images were processed on Olympus Fluoview software v4.2.

### RNA in-situ hybridization (RNA-ISH)

RNA in-situ Hybridization was performed on FFPE mouse tissue using RNAscope^®^ 2.5 HD Duplex Assay kit (ACDBio) following the manufacturer’s instructions. Anti-mouse MCP-1 and CXCL1 probes were used on the tissue slides. mPPIB and dapB were used as positive and negative controls for the tissue samples, respectively. For image analysis, ImageJ 1.52e with Fiji plugin was used to quantify RNA-probe signals.

### 16S rRNA sequencing

Library construction and sequencing was performed at the University of Missouri Metagenomics Core in collaboration with IDEXX BioAnalytics. Bacterial 16S rRNA amplicons were generated using amplification of the V4 hypervariable region and then sequenced using the Illumina MiSeq platform as previously described (31, 32). Paired DNA sequences were merged using FLASH software (33) for a base quality of 31. Cutadapt (https://github.com/marcelm/cutadapt) (34) was used to remove primers and reject contigs that did not contain primer sets. The usearch fastq_filter command (http://drive5.com/usearch/manual/cmd_fastq_filter.html) (35) was used for quality trimming of contigs with rejection of contigs with errors greater than 0.5. Additionally, contigs were clipped to 248 bases with removal of shorter contigs. Output files for samples were concatenated into one file and clustering was performed using the uparse method (http://www.drive5.com/uparse/) (36) for clustering of contigs with 97% similarity and removal of chimeras. Taxonomy was assigned using SILVA database v128 (37).

### Dextran-FITC oral gavage and serum analysis

Intestinal permeability assay was performed according to previously published procedures (38). Briefly, 25-week NZM mice were inoculated with or without segmented filamentous bacteria (SFB) via oral gavage. Two weeks post-inoculation, mice were fasted overnight and received 44mg/100g body weight Dextran-FITC (Sigma 46944) via oral gavage 12h later. Serum was isolated 2 hours post-gavage. Relative intestinal permeability was determined by fluorimeter (excitation 485nm; emission 528nm – Molecular Devices, San Jose, CA).

### Statistical analyses

All statistical analyses for non-microbiota studies used GraphPad Prism 8.0.1 (GraphPad Software, La Jolla, CA, www.graphpad.com). Individual figure legends indicate statistical tests used for different experiments. For GM analysis, bar graphs were generated with Microsoft Excel (Microsoft, Redmond WA) and principal coordinate analysis (PCoA) was generated using Paleontological Statistics Software Package (PAST) 3.12 (39). All groups were visually inspected for descriptive analysis of consistency between samples (bar graphs) or clustering of samples within treatment groups by principal coordinate analysis (PCoA). Statistical testing for differences in alpha-diversity was performed via two-way PERMANOVA, implemented using PAST 3.12. Statistical analysis of differences in family relative and genus relative abundance was performed by one-way ANOVA or Kruskal–Wallis, depending on normality of data as determined via Shapiro-Wilk normality testing, using GraphPad Prism 7.0 for Windows (GraphPad Software, La Jolla, CA, www.graphpad.com). Statistical significance of families or genus were assumed if p□≤□0.05, taking into account false discovery rate using the original method of Benjamini-Hochberg correction for multiple testing (40). Hierarchical clustering and statistical significance at the operational taxonomic unit (OTU) level was determined by two-way ANOVA using MetaboAnalyst 3.0 (41).

## Results

### Exacerbation of destructive kidney disease in mice inoculated with SFB

NZM2410 mice begin to develop glomerulonephritis at approximately 20 weeks of age (42, 43). Glomeruli primarily possess IgG and complement (C3, C4) deposits that eventually glomerular fibrosis and tubulointersitial disease (44). Inflammatory cell invasion predominantly consists of macrophages and dendritic cells and is common to both glomeruli and tubules. To examine the influence of SFB on lupus nephritis in NZM2410 mice we assessed the terminal serum blood urea nitrogen levels (BUN) at 30 weeks of age. Mice colonized with SFB had higher BUN levels than that of SFB-negative mice (Fig. 1A). We further assessed kidney damage by histopathology at the same time point. SFB-negative mice displayed mildly increased cellularity and thickening of glomerular capillary walls upon staining with hematoxylin and eosin (H&E) and Periodic-acid Schiff (PAS) (Fig. 1B). In contrast, mice colonized with SFB possessed markedly enlarged, hypercellular glomeruli with thickened mesangia and hyaline deposits (Fig. 1B). Kidneys of SFB-colonized mice stained with methenamine silver revealed several glomeruli with characteristic subepithelial spike formations on the glomerular basement membrane (GBM). Crescent formation, a hallmark of immune complex glomerulonephritis, was also observed in SFB-colonized mice (data image not shown). H&E and PAS images were reviewed and scored to compare +SFB mice to controls. Overall, glomeruli from +SFB mice had more severe glomerulonephritis and were identified as being larger, more proliferative, possessing more hyaline deposits as well as cellular crescents than -SFB controls (Fig. 1C). Although +SFB and -SFB mice both possessed glomerular abnormalities, only +SFB mice had aberrations in their tubulointerstitium as manifested by tubular protein casts, tubular degeneration and necrosis with pronounced tubular regeneration, and mild interstitial inflammation (Fig. 1C). IgG and C3 deposition was evident in +SFB and -SFB mice. However, the intensity of staining in both cases was significantly more prominent in mice inoculated with SFB (Fig. 1D).

**Figure 1.**
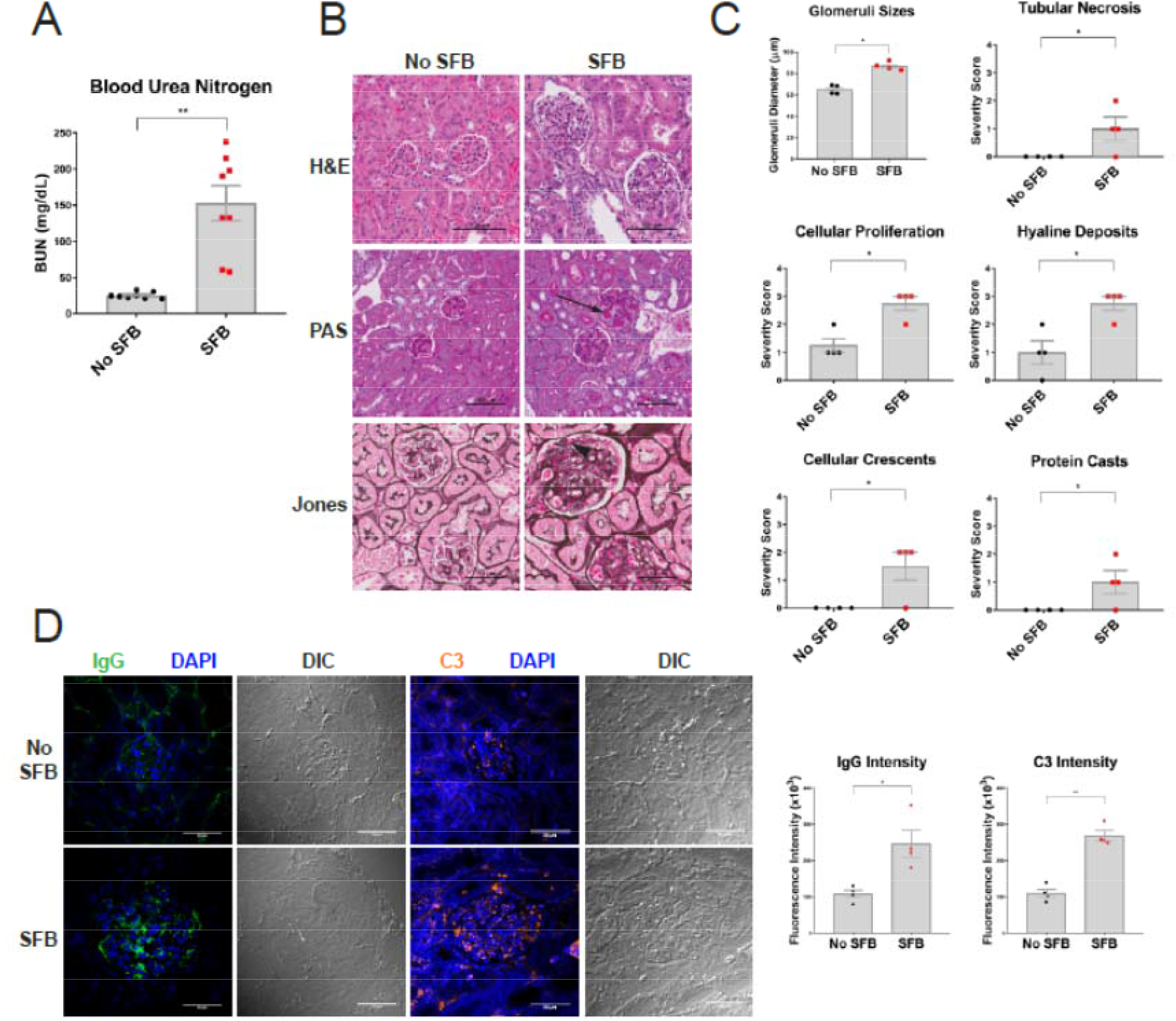
NZM2410 colonized with SFB exhibit intensified kidney disease with elevated immune-complex deposition. 10-week old mice were oral gavaged with fecal matter from mice harboring SFB or control mice and sacrificed at 30-weeks of age and subjected to biochemical analysis. **A.** Serum blood urea nitrogen (BUN). **B.** Kidney disease was directly assessed by performing hematoxylin & eosin (H&E), periodic acid-Schiff (PAS) and Jones’ silver stains. H&E, PAS and silver stain highlight enlarged glomeruli, hyaline deposits (black arrow) and subendothelial deposits (black arrowhead) in SFB colonized mice, respectively. **C.** Histopathological assessment of kidney tissue was blindly scored by a veterinary pathologist; kidney pathology was defined as tubular necrosis, cellular proliferation, hyaline deposits, cellular crescents and protein casts. **D.** Immunofluorescence staining of IgG and C3 deposition in the glomerular and tubulointerstitium of kidney tissue; differential interference contrast (DIC) also shown. Error bars represent mean ± SEM (A, C and D). Unpaired Student t test (A, C and D). *p<0.05; **p<0.005.

### SFB inoculated mice produce elevated inflammatory cytokines and chemokines

The role of cytokines in the disease pathogenesis of LN is not fully understood. However, overwhelming evidence suggests that particular proinflammatory cytokines and chemokines are elevated in lupus (45–48). Previous studies have shown that the chemokines MCP-1, MIP-1α and MIP-1β are increased in the serum of patients with SLE (49, 50). Patients with LN have been shown to have elevated urinary MCP-1 that correlates with disease activity (46). In mice, knocking out or neutralizing MCP-1 or macrophage-derived chemokine (MDC), respectively, led to a decrease in the macrophage infiltration into the kidney as well as a reduction in glomerulonephritis (47, 51). To investigate the cytokine profile in NZM2410 mice colonized with SFB we analyzed serum at 30 weeks of age. The proinflammatory cytokines IL-6 and IL-17A were significantly elevated (Fig. 2A). The prototypical proinflammatory cytokines IL-1β and TNF-α were elevated in +SFB mice, although not significant (Fig. 2A). The macrophage chemoattractants MCP-1 and CXCL1 were increased in the serum of +SFB mice at 30 weeks of age (Fig. 2A). We then tested for gene and protein expression of these chemokines in the kidney using RNAScope and IHC, respectively. We found that MCP-1 and CXCL1 expression is significantly increased in the kidneys of +SFB mice at the same time point (Fig. 2B and C). Both glomeruli and tubules were involved. However, chemokine expression tended be periglomerular and/or tubular rather than intraglomerular (Fig. 2B and C).

**Figure 2.**
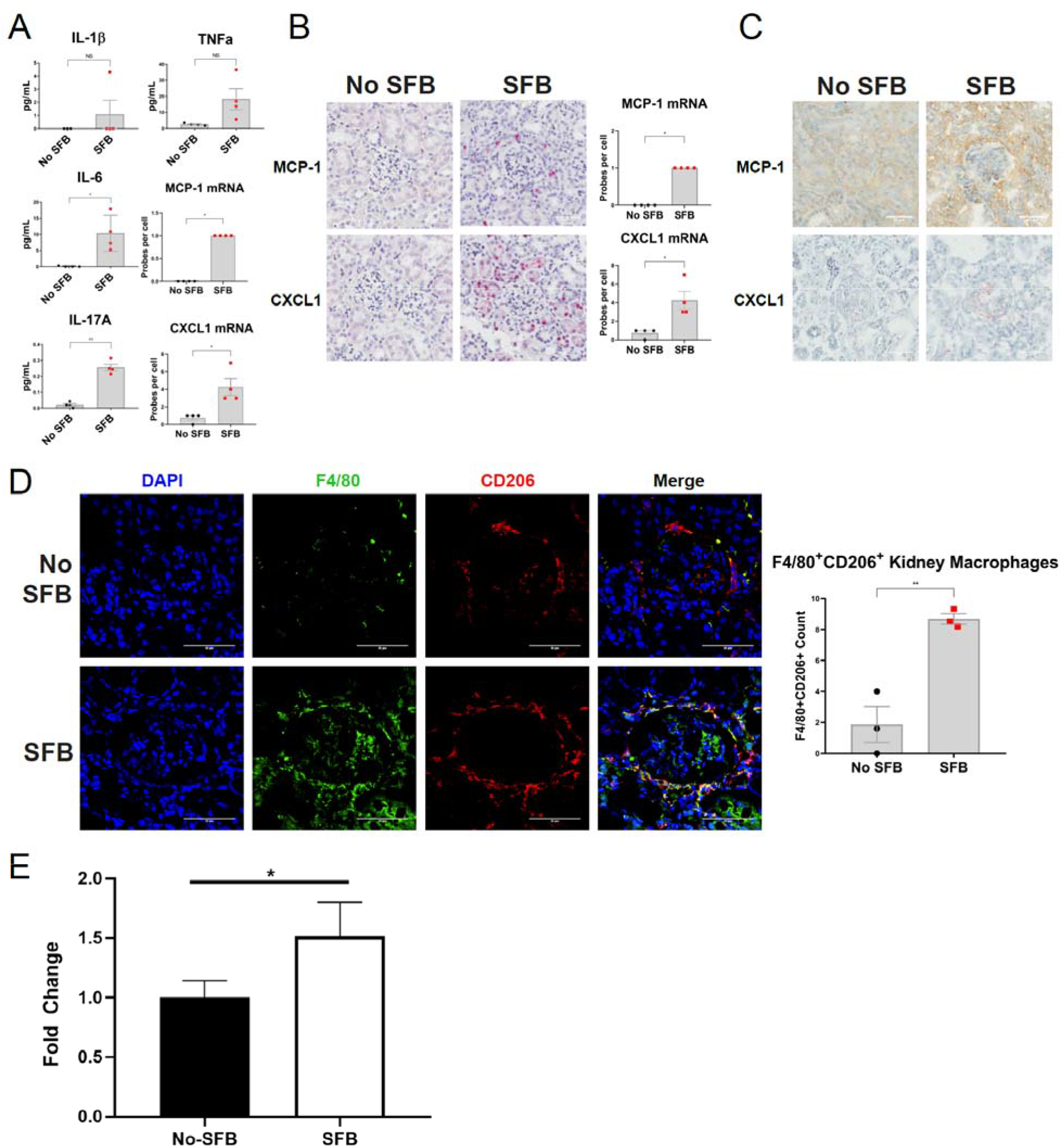
Macrophage chemoattractants and M2-like macrophages are elevated in the kidneys of NZM2410 mice colonized with SFB. **A**. Proinflammatory serum cytokines and chemokines as measured by ELISA at 30 weeks of age. **B**. mRNA levels of MCP-1 and CXCL1 in kidney tissue as measured by RNAScope. **C**. IHC staining for MCP-1 (DAB staining) and CXCL1 (AEC staining) proteins in kidney tissue. **D**. Immunofluorescence staining of kidney tissue for F4/80 (green), CD206 (red) and nuclei (blue). **E**. Intestinal permeability is increased in SFB-colonized mice as measured by Dextran-FITC fluorescence;. Error bars represent mean ± SEM (A, B, D). Unpaired Student t test (A, B, D). NS, not significant; *p<0.05; **p<0.005.

### Increased CD206+ macrophage kidney infiltration into SFB colonized mice

The disease pathogenesis of lupus nephritis is complex and the cellular component that drives these inflammatory processes remains unresolved. Renal infiltrating cells in the kidneys of lupus patients and lupus mouse models predominately consist of macrophages and T cells (51–56). Multiple T cells subtypes within the CD4 and CD8 lineage as wells as double negative T cells have been observed in the kidneys of patients and mouse models with LN (53, 57, 58). Since SFB is known to induce Th17 cell differentiation and proliferation we stained kidney tissue from 30-week old +SFB and -SFB NZM2410 mice for CD4 and RORγt. Although we observed an increase in Th17 cells in the small intestinal lamina propria of SFB exposed mice (Fig. 3A), we did not detect CD4+RORγt+ cells in either cohort of mice (data not shown). However, given that the macrophage chemokines MCP-1 and CXCL1 were elevated in the serum as well as kidney tissue of +SFB mice we tested for the presence of macrophages in the kidneys of these mice and controls. Infiltrating F4/80+ myeloid cells were observed circumscribing the glomeruli in both no SFB and SFB cohorts (Fig. 2D). To further delineate the subtype of macrophage we stained kidney tissues for F4/80, CD68 and CD206. F4/80+CD68+ macrophages were not detected in either +SFB or control mice (data not shown). However, many of the kidney F4/80+ macrophages also costained for CD206 and the number of F4/80+CD206+ macrophages was also greater in mice SFB colonized mice (Fig. 2D).

**Figure 3.**
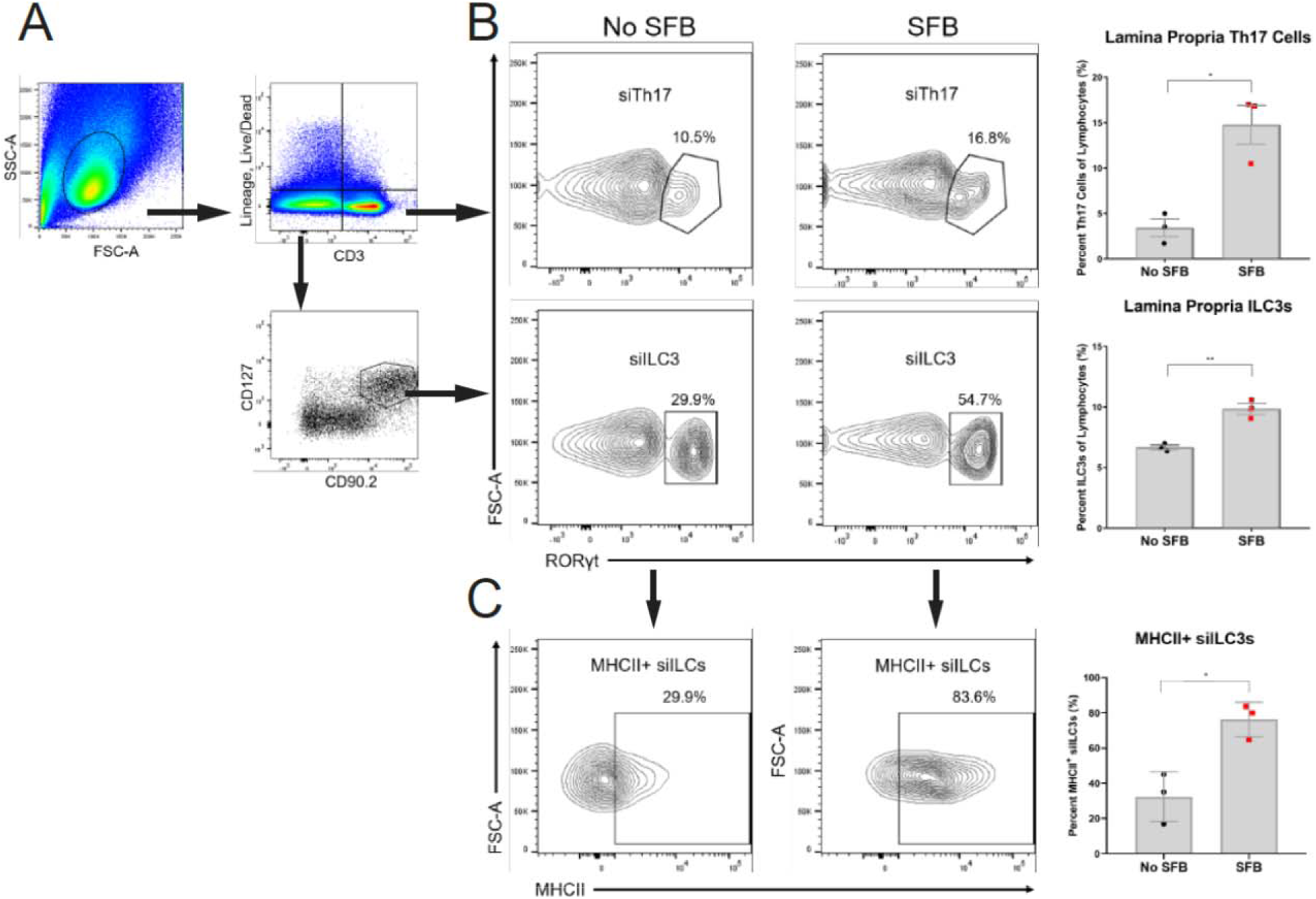
Expansion of small intestinal Th17 cells and ILC3s in SFB colonized NZM2410 mice. **A**. Gating strategy of small intestinal lamina propria Th17 (siTh17) cells and ILC3s (siILC3) in 30 week old mice. Gating is based on previous gating on the lymphocyte gate (top left panel), live and lineage negative markers (CD1a, CD3, CD4, CD14, CD11b, CD19, NK-1.1, F4/80). **B**. RORγt+ siTh17 cells and siILC3s in mice colonized with SFB when compared to control mice (No SFB). **C**. MHCII expression in siILC3s. Error bars represent mean ± SEM (B and C). Unpaired Student t test (B and C). *p<0.05; **p<0.005.

### SFB induced expansion of Th17 cells and ILC3s

SFB induces intestinal expansion of Th17 cells and ILC3s in GF and SPF wildtype mice (22). Although Th17 cell and ILC3 expansion incurs no observable pathogenic effect on wildtype mice, IL-17+ T cells have been correlated with disease in LN. Contrastingly, the role of ILC3s in SLE and/or LN is not known. However, ILC3s are capable of both regulatory and pathogenic functions in other autoimmune diseases (59, 60). Moreover, a key relationship between ILC3s and the microbiota is thought to exist and is in part maintained by MHCII interactions (61). Therefore, we were interested in testing whether +SFB NZM2410 mice with worse kidney disease had expanded Th17 cell and ILC3s than -SFB mice. Serum IL-17A was increased in 30-week old SFB-colonized mice when compared to controls (Fig. 2A). However, kidney expression of IL-17A and IL-22 was undetectable in either +SFB or -SFB mice (data not shown). We then tested whether the small intestine experienced a local expansion of Th17 cells and/or ILC3s (siTh17 and si ILC3s, respectively) as was reported in wildtype mice colonized with SFB (Fig. 3A) (22). Mice colonized with SFB had an expansion of both siTh17 cells and siILC3s (Fig. 3B). We also looked more closely into the phenotype of the siILC3s and found that these ILC3s express higher levels of MHCII, suggesting a higher potential for these cells to antigen present (Fig. 3C). To test for renal infiltration we performed IHC on kidney tissue. As reported in previous literature (53, 58) we observed intraglomerular and periglomerular CD3+ T cells in both +SFB and -SFB mice, however, we did not detect infiltrating RORγt+ Th17 or ILC3s in the renal parenchyma (data not shown).

### Reduced intestinal tight junction expression with SFB colonization

The small intestine is the primary site of nutrient absorption from ingested food and this function is evident in its relatively large surface area. Like the skin, the small intestinal mucosa also acts as a barrier between the outside environment and host. The large surface area presents a surveillance challenge for the immune system. One of the ways to overcome this issue is the placement of tight junctions along the lateral membrane of intestinal epithelial cells. The claudin and ZO family of tight junction proteins ensure proper sealing between two epithelial cells (62, 63). Defects in either family contribute to increased intestinal permeability, also known as “leakiness”. Gut leakiness has been associated with autoimmunity wherein the microbiota is also perturbed (13). Therefore, we investigated the tight junction integrity between SFB colonized and non-colonized NZM2410 mice. Among 30 week old SFB inoculated mice, expression of the tight junction scaffolding protein ZO-1 was reduced (Fig. 4A). Similarly, claudin-1 and claudin-3 expression was reduced when compared to mice without SFB, suggesting an overall decrease in intestinal barrier integrity (Fig. 4B).

**Figure 4.**
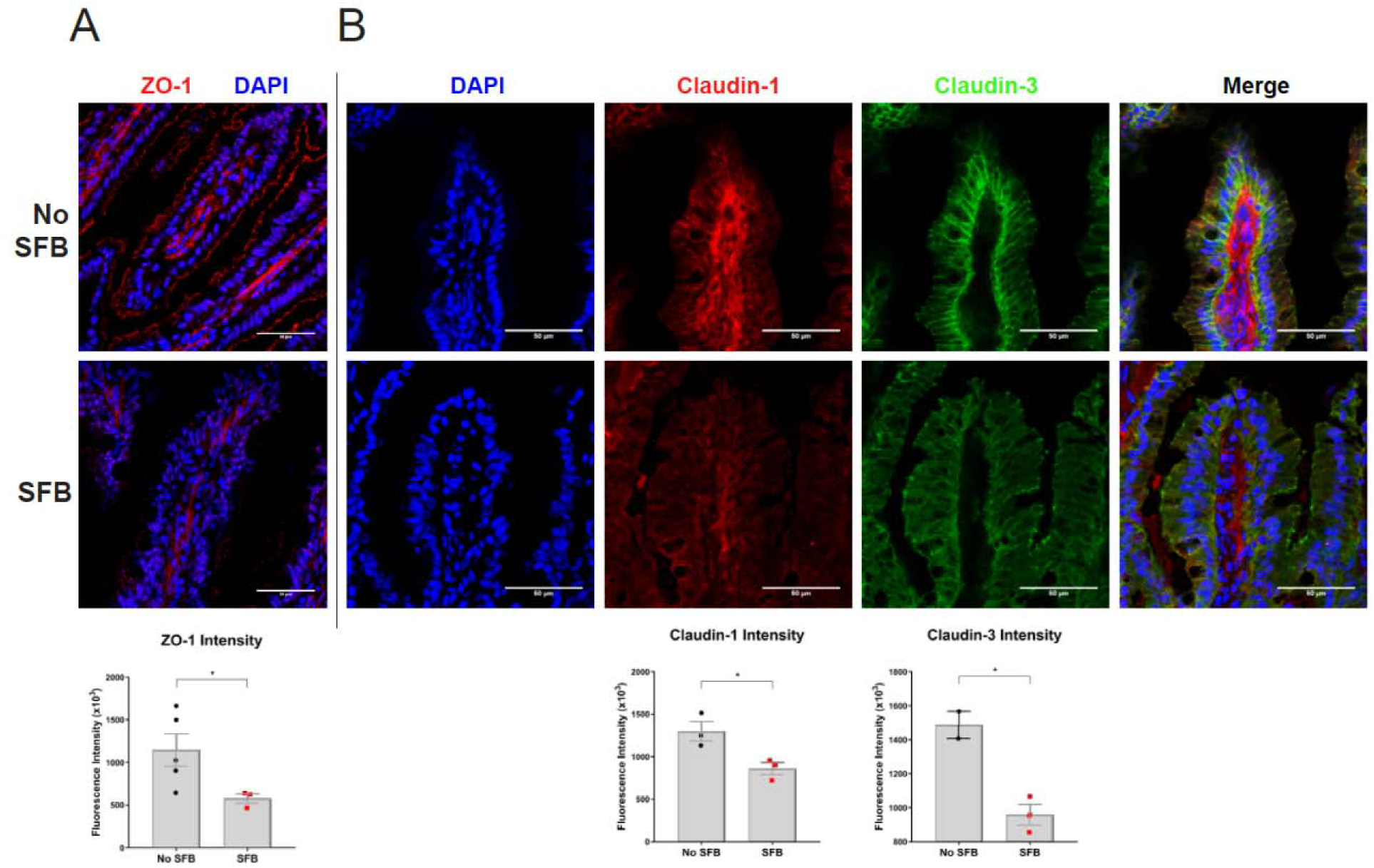
Small intestinal tight junctions are reduced in SFB colonized NZM2410 mice. **A**. ZO-1 tight junction scaffolding protein (red) is decreased in +SFB mice at 30 weeks of age. **B.** Claudin-1 (bottom panels; red) and Claudin-3 (bottom panels; green) tight junctions were also decreased in +SFB mice at the same time point. Standardized fluorescence intensity analyses are shown below. Error bars represent mean ± SEM (A and B). Unpaired Student t test (A and B). *p<0.05.

### Intestinal permeability increases with SFB colonization

One method of assessing intestinal permeability is by use of an orally administered Dextran-FITC conjugate that is subsequently absorbed by the small intestine and then quantified in the serum (38). We tested the serum of 25-week old Dextran-FITC gavaged NZM2410 mice that were either SFB colonized or not. Following oral gavage, we found that +SFB mice had significantly elevated Dextran-FITC concentrations in their serum when compared to controls. Furthermore, these findings were observed at both 1 and 2 hour post-gavage time points (post 1 hour data not shown).

### Intestinal GM dysbiosis is influenced by the presence of SFB

A limited number of lupus-specific GM studies have been performed and even less have investigated LN (11–16). Some of these studies observed differences in particular GM phylogenetic taxa between healthy individuals and patients with SLE (14–16). Studies have also investigated the overall changes in the intestinal microbiota in various mouse models of lupus (11–13, 16). However, to our knowledge only one study has specifically investigated the role of the GM in LN (13). This study identified a beneficial bacterium within Lactobacillus. To provide an in-depth impression of the intestinal bacterial microbiome we utilized 16S ribosomal RNA (rRNA) sequencing to detect unique hypervariable regions that correspond to phylogenetic markers in 16S rRNA (31, 32). Fecal DNA was purified from mice at either 15 or 30 weeks of age. Overall, 529 taxa were detected across all samples. Sequencing depth and calculation of relative abundance allowed for analysis down to the species-level in certain instances (Fig. 5). Most of the represented operational taxonomic units (OTUs) have not been cultured, isolated and sequenced at the species level and can only be classified up to the Genus or Family taxa. We were unable to classify Clostridiaceae OTUs that matched with SFB. However, SFB was detectable by using SFB-specific 16S ribosomal DNA (rDNA) primers during PCR of fecal DNA (Supplemental Fig. 1A and 1B) (30). Relative abundance OTU differences were detected across all four cohorts of mice: at 15 or 30 weeks of age and +/- SFB (Fig. 5).

**Figure 5.**
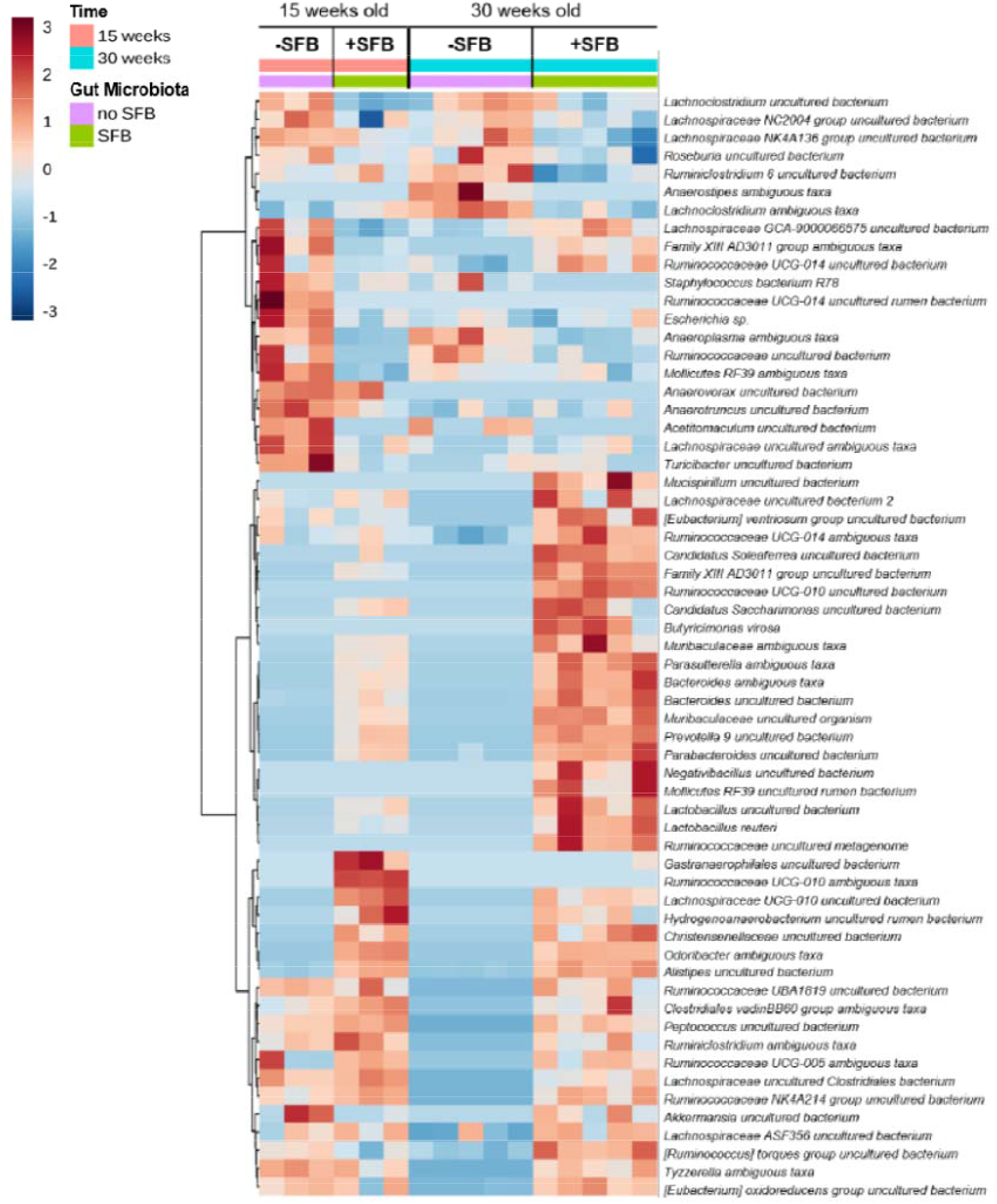
Hierarchical cluster analysis of statistically significant operational taxonomic units (OTUs) in each group based on the presence of SFB and time. Color intensity indicates Log2 normalized abundance of OTUs in each sample. Color coded bars at top indicate SFB status and time of sample collection.

Principal coordinate analysis (PCA) revealed four distinct clusters across three axes that corresponded to no SFB at 15 weeks old, +SFB at 15 weeks old, no SFB at 30 weeks old and +SFB at 30 weeks old (Fig. 6A). The greatest variability was identified in the PC1 axis and separated the no SFB and +SFB groups. Principal component analysis (PCoA) also showed that the greatest variability was between these two groups (Supplemental Fig. 2A). Statistical significance was achieved up until the 5^th^ PCoA (data not shown). In addition, separate clustering was observed between the +SFB 15 and 30 week old cohorts, suggesting that an important change in the intestinal microbiota occurred after the onset of disease.

**Figure 6.**
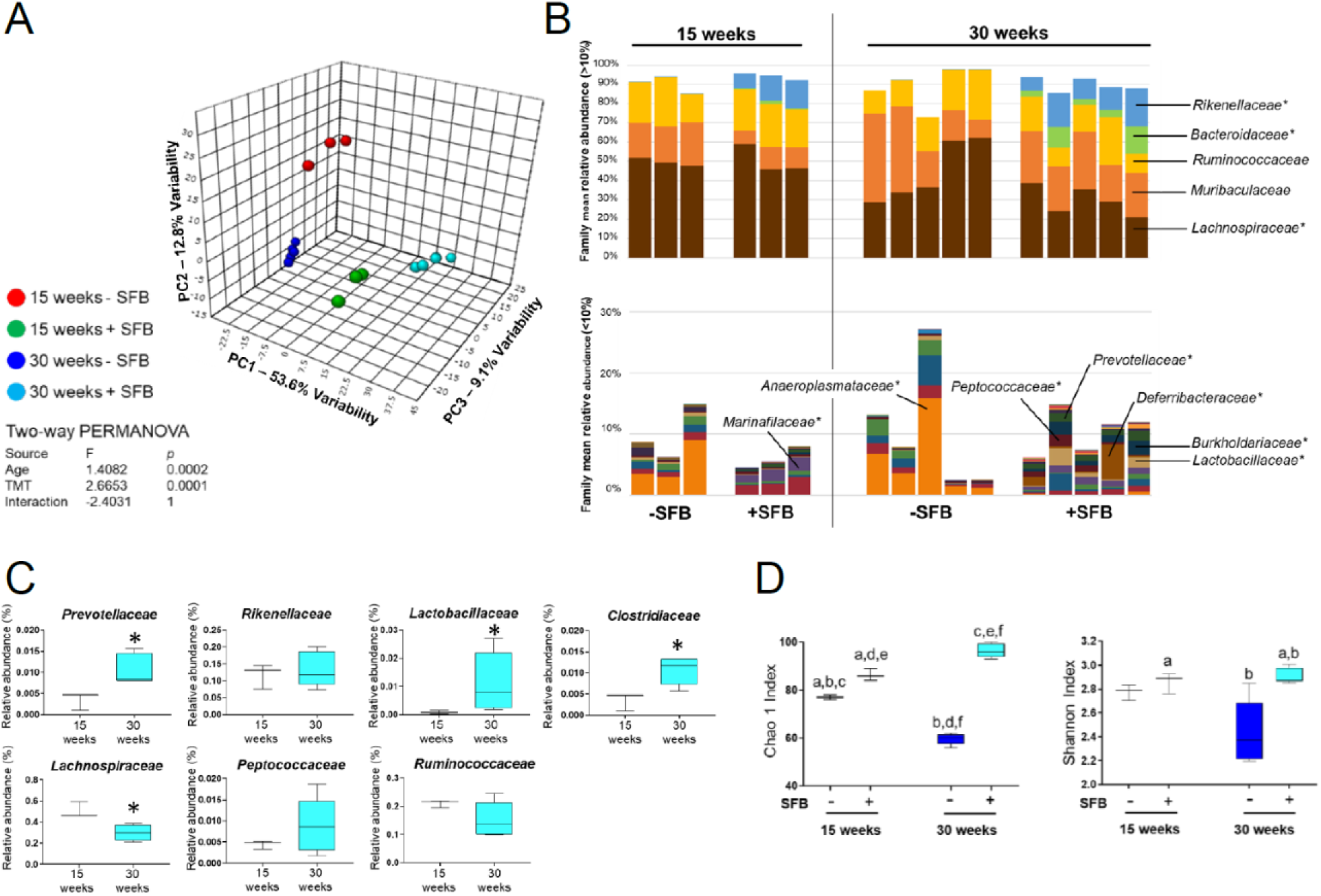
Gut bacterial microbiome changes between +/-SFB NZM2410 mice at 15 or 30 weeks of age. **A**. Principal coordinate analysis (PCoA) of representative fecal samples from SFB negative or SFB positive colonized mice at 15 and 30 weeks of age. Figure legend of groups located at right of PCoA. Statistical significance determined using two-way PERMANOVA (p≤0.05 statistically significant). **B**. Bar charts of relative abundance of taxa at family level from SFB negative or SFB positive colonized mice at 15 and 30 weeks of age. Statistical significance determined using two-way ANOVA or Kruskal-Wallis, depending on normality of data as determined by Shapiro-Wilk normality testing and Benjamini-Hochberg correction for multiple testing (p≤0.05 statistically significant). Statistically significant families indicated by asterisk. **C**. Percent relative abundance of significant taxa at the family level from representative fecal samples of SFB positive mice at 15 and 30 weeks of age. Statistical significance determined using the Mann Whitney test (p≤0.05 statistically significant). Asterisks denote statistical significance between groups. **D**. Chao1 and Shannon estimate of microbial richness and diversity plotted by Tukey box and whisker graph (n=3-5 per group). Statistical significance determined using two-way ANOVA (p≤0.05 statistically significant). Statistical significance between groups annotated by same lower case letters above plots.

Similar to the Genera level differences observed (Fig. 5), compositional changes were detected at the Family level in lupus mice SFB colonized at 15 and 30 weeks of age (Fig. 6B). Expanded Families included Prevotellaceae, Lactobacillaceae and Clostridiaceae. We also detected reduced relative abundance of Ruminococcaceae and Lachnospiraceae (Fig. 6B and 6C). When comparing -SFB to +SFB mice at 30 weeks we observed significant increases in Prevotellaceae, Rikenellaceae, Lactobacillaceae and Peptococcaceae (Supplemental Fig. 2B). Clostridiaceae, Ruminococcaceae and Lachnospiraceae, however, did not experience a significant change in relative abundance at 30 weeks with or without SFB present.

To assess GM biodiversity in the presence or absence of SFB as well as during the progression of lupus nephritis we calculated the alpha diversity contained within the 16S rRNA. Alpha diversity encompasses the Chao 1 and Shannon indexes wherein Chao 1 estimates total species richness by assigning weight to rare OTUs and Shannon takes into account abundance and evenness within a microbial community and is also sensitive to rare OTUs (64, 65). The Chao 1 alpha diversity was significantly greater in both the +SFB 15 and 30 week old mice when compared to no SFB cohort of the same ages (Fig. 6D). Chao 1 index decreased in the no SFB group at 30 weeks of age when compared to the same group at 15 weeks of age. However, the opposite effect was observed in the +SFB group at 30 weeks of age when compared to the same group at 15 weeks of age. Similar trends are observed with respect to the Shannon index, suggesting that possessing SFB enhances intestinal biodiversity the bacterial community. Altogether, SFB containing mice exhibit higher intestinal biodiversity than the no SFB group.

Given a recent publication that focused on GM dysbiosis in LN patients (66), we were interested in revisiting our dataset to find any potential similarities. In this report, *Ruminococcus gnavus* was found to be elevated in LN patients as compared to healthy subjects (66). While we could not detect the reported LN candidate pathobiont *R. gnavus*, we were able to detect a group of relatives, *Ruminococcus torques* spp. In our dataset, we found that 30 week old SFB+ mice had a significant enrichment of *R. torques* spp. (also known as *R. torques* group) when compared to 15 week old SFB+ mice which advocates that perhaps disease severity was an important factor in this change (Supplemental Fig. 3A and 3B). Additionally, 30 week old SFB+ mice had greater relative abundance of *R. torques* spp. when compared to 30 week old SFB-mice, suggesting the potential of an SFB influence. Interestingly, we could not detect *R. torques* spp. in either SFB+ or SFB-B6 WT mice at 15 or 30 weeks of age (data not shown). We then used PATRIC, a phylogeny bioinformatics database to compare the relationship between *R. torques* spp. and *R. gnavus* (67). Cladogram analysis revealed that *R. torques* spp. and *R. gnavus* shared a common ancestor back four species and that within the genera *Ruminococcus, R. torques* spp. and *R. gnavus* were most closely related suggesting that these bacteria are highly related (Supplemental Fig. 3B). Our analysis is supported by previously published 16S rDNA sequencing analysis (68).

## Discussion

The steady state relationship between the host and commensal organisms functions to maintain host access to nutritional resources such as essential vitamins and short-chain fatty acids (27). Furthermore, the influence of the microbiota on the host immune system is now widely accepted that dysbiosis can lead to human disease (28). One of the first studies to examine the role of the microbiota in autoimmunity observed that germ-free NOD mice developed more severe autoimmune diabetes when compared to SPF-raised NOD mice (69). Recent reports have uncovered similar findings in human autoimmune diseases (6–8). A small number of studies have specifically investigated the intestinal microbiota’s role in human and murine lupus (11, 13–16). In this study we were interested in the effects of SFB on lupus phenotype and specifically kidney disease in NZM2410 mice. Wildtype mice colonized with SFB have no discernable health deficits (22). However, SFB induces more severe disease in inflammatory arthritis models (17, 26–28). Our results demonstrate that glomerulonephritis severity, immune cell aberrations and intestinal barrier abnormalities are all influenced by SFB colonization in our LN model. Moreover, the GM is substantially altered with the presence of SFB.

Kidney infiltrating immune cells are considered to be important in the pathogenesis of LN (51–54). Activated CD4 and CD8 T cells and their effector functions are hypothesized to be the key players in immune-mediated kidney damage in LN (70, 71). Given that SFB gut colonization induces small intestinal Th17 cell proliferation in wildtype mice (22) as well as in our study (Fig. 3B) we were interested in investigating the role that SFB might play in promoting Th17-associated kidney damage. Similar to observations reported in other diseases (53, 72), we observed T cell infiltrates in the kidneys of +/-SFB mice (data not shown). However, in our study we did not detect kidney infiltrating CD4+RORγt+ Th17 cells in either +SFB or -SFB NZM2410 mice at 30 weeks of age (data not shown). This finding may be in part due to recent evidence that reports kidney infiltrating CD4+ or CD8+ T cells are not effector cells in LN and in fact maintain an “exhausted” transcriptional as well as functional profile (72). Although correlations between LN severity and abundance of CD4+IL-17+ Th17 cells has previously been made, this study suggests that the pathogenic importance of these and other T cell subtypes in LN is limited. Interestingly, coupled with our findings that show increased SFB positivity in NZM2410 mice is linked to SILP Th17 cells and ILC3, questions about the significance of this result arise. Increased ILC3 MHC Class II expression is an especially intriguing finding as this mirrors what was found in Crohn’s disease patients. MHCII+ ILC3s are known to help control T cell responses through Ag presentation (73). Although we know that mice colonized with SFB have SILP Th17 cells that recognize SFB (24), it is unclear what the significance of ILC3 MHCII expression is on disease states such as LN. Use of ILC3-specific MHCII KO mice or MHCII neutralizing Abs in Rag KO mice would be useful to delineate the mechanism and significance of this finding. Importantly, these transgenic alterations would have to be incorporated into a lupus model background. Therefore, more work is needed to understand this complex relationship.

We demonstrated that the macrophage chemokines MCP-1, MIP1-alpha and CXCL1 expression were elevated in sera of +SFB mice (Fig. 2B and 2C). Moreover, we observed a greater number of F4/80+ macrophages in +SFB mice (Fig. 2D). We further distinguished between M1- and M2-like kidney infiltrating macrophages by staining for CD68 (data not shown) or CD206, respectively. We detected periglomerular CD206+ macrophage staining. Furthermore, these F4/80+CD206+ cells were almost exclusively found in +SFB mice (Fig. 2D). Although M2-like macrophages were traditionally considered to be “anti-inflammatory” data suggests functional plasticity (74). Studies have shown that M2-like macrophages are involved in the pathogenesis of LN (55, 56). CD206 is a scavenger receptor capable of clearing autoantigens (75). Interestingly, deletion of CD206 in an immune-complex glomerulonephritis model led to a significant reduction in kidney disease. Furthermore, macrophages from CD206 knockout mice exhibited reduced phagocytic capacity when exposed to apoptotic kidney mesangial cells (55). In humans, CD206+ kidney infiltrating macrophages were associated with more active disease (56). These particular cells were also more numerous surrounding the glomeruli of LN patients than other forms of glomerulonephritis like IgA nephropathy and the ANCA-associated vasculitides (AAV). The role that CD206+ macrophages play in gut dysbiosis in LN is unknown. It is possible that these infiltrating cells are activated by SFB and subsequently traffic to the kidney. It would be important to investigate whether intensified kidney disease is in fact carried out by these cells and importantly, mediated by SFB.

Previous studies involving lupus mouse models have found evidence of intestinal dysbiosis (11–13, 16). However, to our knowledge none have identified a commensal pathobiont in LN. Presence of SFB was associated with gut dysbiosis at early and late disease. During more severe glomerulonephritis stages (30 weeks of age), +SFB mice had significantly higher abundances of bacterial Families like *Prevotellaceae* and *Rikenellaceae*. +SFB mice had reduced *Lachnospiraceae* abundance later in disease at 30 versus at 15 weeks. Studies on human lupus GM dysbiosis observed similar changes in the microbiota with disease (14, 15). When looking at higher taxonomic levels, the two most dominant bacterial phyla within the mouse and human GM, Firmicutes and Bacteroidetes, exhibit reduced Firmicutes/Bacteroidetes ratio in patients with SLE when compared to healthy controls (14, 15). In the present study, although not statistically significant we observed a similar trend in +SFB mice at 30 weeks of age (Supplemental Fig. 2C). Relative outgrowth and depletion of certain members of the microbiota may be of importance to overall disease state (76). Therefore, a more in depth look into perturbed bacterial species is required. An interesting note is that the murine lupus model microbial biodiversity defined in the literature does not replicate what is found in the human GM of SLE patients (11, 13–15). Despite lower GM diversity in patients with SLE, greater biodiversity in mice with worse disease has been observed in mouse models of lupus.

While SFB may not be a causative agent of autoimmunity, a recent human study showed that certain species may act as a primer of autoimmunity (66). Similar to our findings, the authors of this particular human study showed that LN patients had dysbiotic GMs. GM dysbiosis also correlated with disease activity. They showed that the species *Ruminococcus gnavus* was the most pronounced in relative abundance in patients with LN. Intriguingly, fecal and serum samples from LN patients had high IgA and IgG that was specific for *R. gnavus*. Furthermore, anti-dsDNA autoantibody from LN patients strongly reacted with *R. gnavus* antigens. We demonstrated that 30 week old +SFB mice had reduced Lachnospiraceae abundance when compared to 15 week old +SFB mice. However, when analyzing the genus level, we only could detect *Ruminococcus torques* spp. Our data showed that the *R. torques* group was significantly enriched in 30 week old SFB+ NZM2410 mice. We searched the PATRIC phylogeny database and found that *R. gnavus* and the *R. torques* were highly related (67). Remarkably, both *R. gnavus* and the *R. torques* are known mucolytic commensal bacteria (77). Indeed, we showed in chapter 2 that SFB positivity was associated with downregulation of tight junction proteins in the small intestinal epithelium. These findings may be a mechanism by which commensals or their antigens might escape the lumen to activate host immune cells. Coupled with the fact that these bacteria are elevated in LN and that serum anti-dsDNA autoAb from LN patients was reacted with R. gnavus Ags suggests that R. gnavus is a potential pathobiont in SFB. More work is needed, however, to test whether R. gnavus alone is capable of worsening kidney disease in GF LN mice as well as in mice raised in a specific pathogen free (SPF) environment.

Collectively, our study demonstrated that changes in GM were associated with detrimental effects to the kidney in a lupus mouse model. Colonization of these mice with SFB greatly enhances inflammatory kidney lesions as suggested by both immune complex deposition as well as immune cell infiltration. Evidence of intestinal barrier aberrances in mice colonized with SFB was noted as well and suggests a role for pathobionts with this capacity. Moreover, disease severity was associated with gut dysbiosis. Given recent data, specific species such as *R. gnavus* may represent primers of autoimmunity and therefore warrant further investigation. Future studies are needed to further clarify the relationship between the influence that SFB may have on altering the host immune system as well as the rest of the intestinal microbiota.

## Notes

### Competing Interest Statement

The authors have declared no competing interest.

