## Supplemental Figures for "Gut Dysbiosis with a Pathobiont Shifts the Intestinal Microbiota Profile and Accelerates Lupus Nephritis"

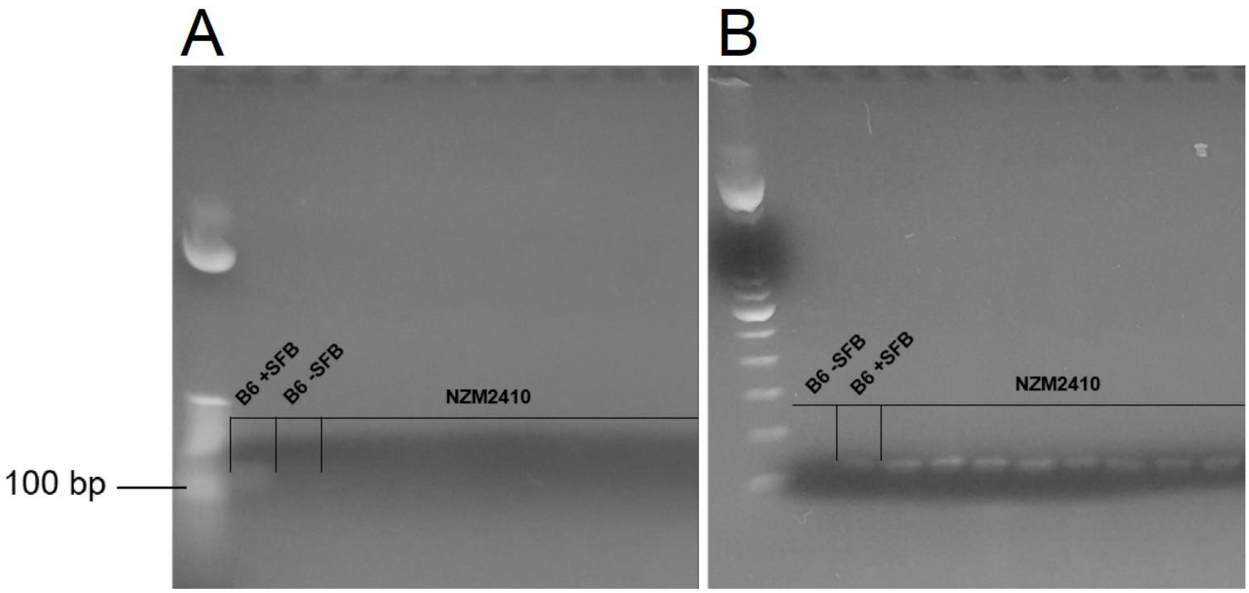


**Supplemental Figure 1**. SFB testing in NZM2410 mice inoculated with B6 -SFB or B6 +SFB fecal homogenates. SFB-specific 16S rDNA primers by PCR is shown at 108 base pairs. NZM2410 mice were tested at 30 weeks of age after being oral gavaged with B6 –SFB **A**. or B6 +SFB **B**. fecal homogenates at 10 weeks of age. B6 +/- SFB mice were purchased from Taconic Farms; mice were tested by the vendor and upon arrival to our vivarium for SFB.

A

Two-way PERMANOVA

| Source | Sum of sqrs | df | Mean square | F | p |
|---|---|---|---|---|---|
| Age | 0.68251 | 3 | 0.2275 | 1.4082 | 0.0002 |
| TMT | 0.43059 | 1 | 0.43059 | 2.6653 | 0.0001 |
| Interaction | -1.1647 | 3 | -0.38822 | -2.4031 | 1 |
| Residual | 1.2924 | 8 | 0.16155 | | |
| Total | 1.2409 | 15 | | | |

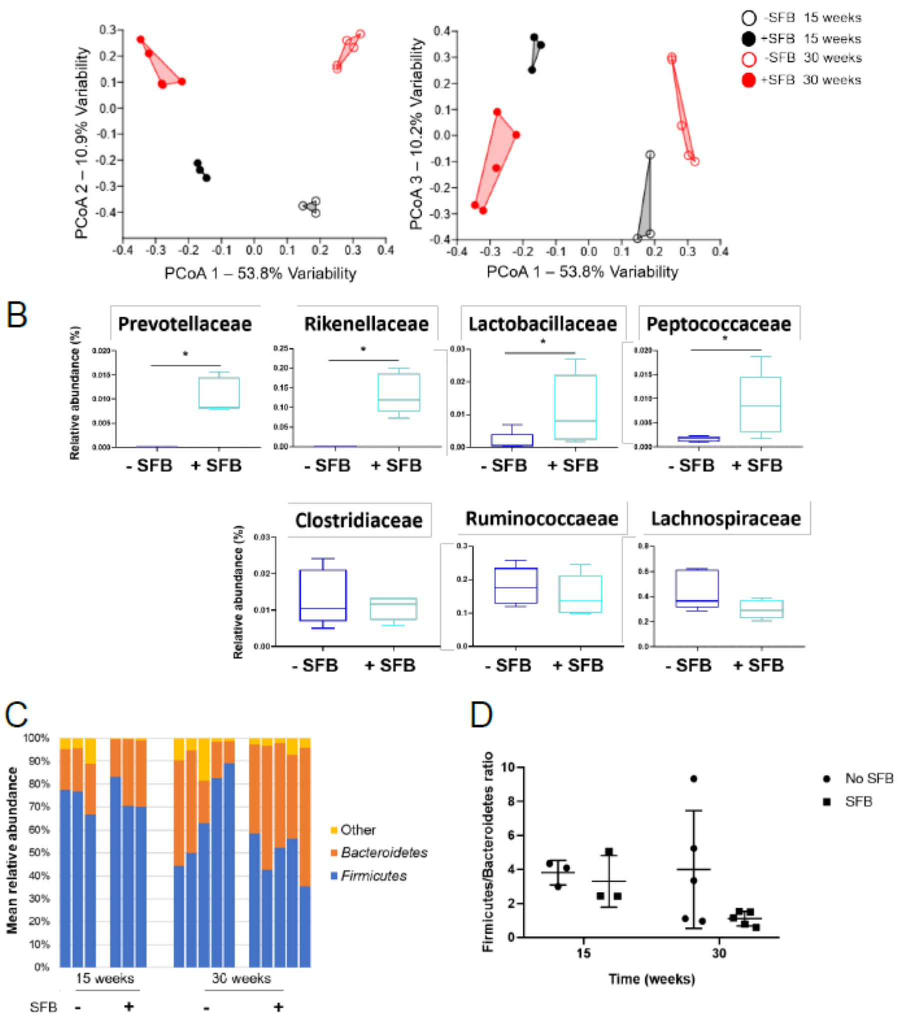


**Supplemental Figure 2**. 16S rRNA analysis of NZM2410 mice +/-SFB at 15 or 30 weeks of age. **A**. Principal Component Analysis (PCoA) of all operational taxonomic units (OTUs) in the four cohorts of mice. The bulk of the variability in the groups is noted in PCoA1 and PCoA2. **B**. Percent relative abundance of significant taxa at the family level from representative fecal samples of SFB positive mice at 30 weeks of age. Statistical significance determined using the Mann Whitney test ($p \leq 0.05$ statistically significant). Asterisks denote statistical significance between groups. **C**. Bar charts of relative abundance of taxa at the Firmicutes and Bacteroidetes Phylum level at different 15 and 30 weeks, with and without SFB. **D**. Firmicutes to Bacteroidetes ratio between the same cohorts in A. Statistical significance was not achieved as determined by two-way ANOVA or Kruskal-Wallis, depending on normality of data as determined by Shapiro-Wilk normality testing and Benjamini-Hochberg correction for multiple testing ($p \leq 0.05$ statistically significant).

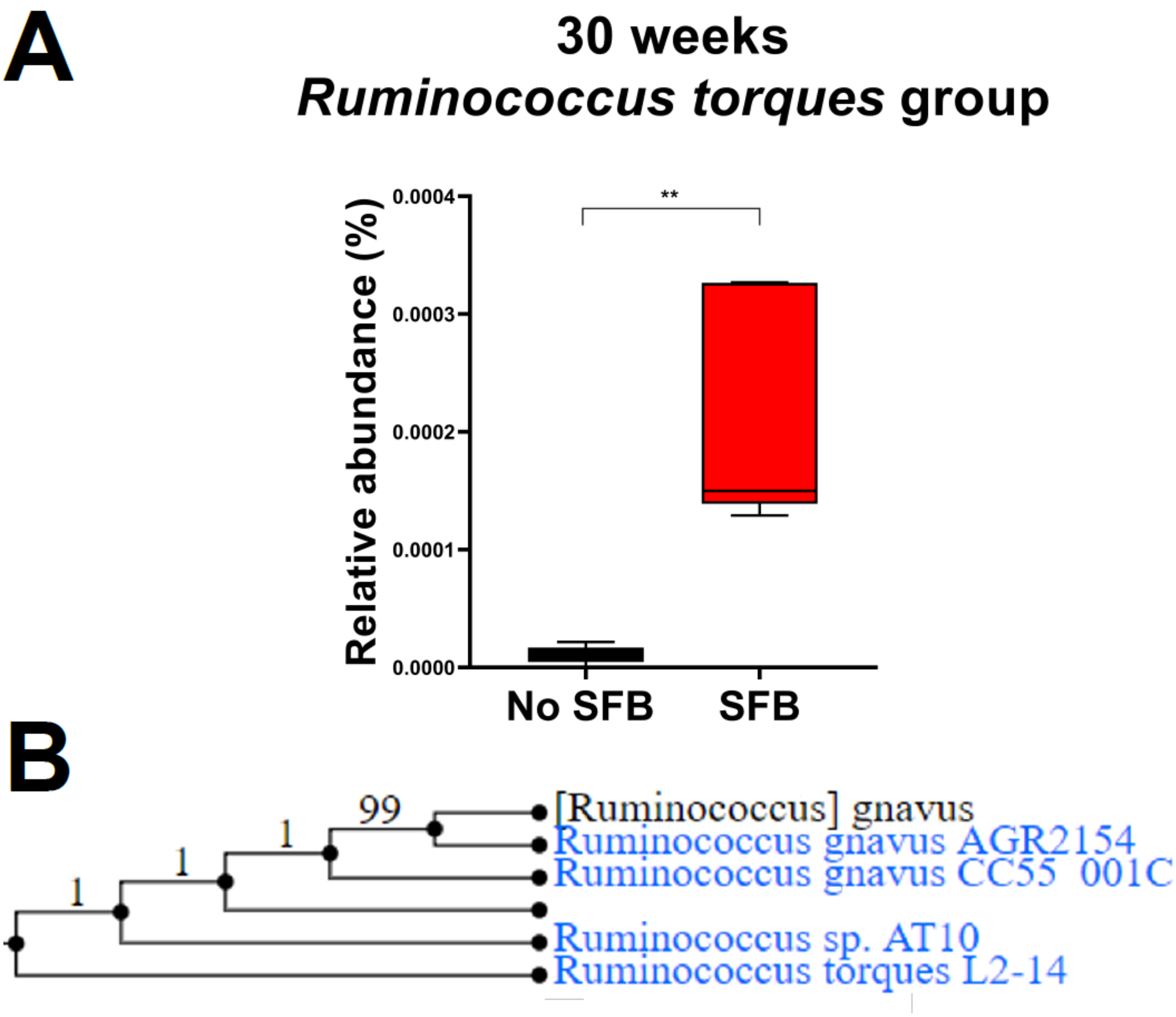


**Supplemental Figure 3**. *Ruminococcus torques* group from 16S rDNA analysis of +SFB and –SFB NZM2410 mice at 30 weeks of age (n = 3-5 mice per group). **A**. Comparison of *R. torques* species in +SFB and −SFB NZM2410 mice at 30 weeks of age. **B**. Phylogenetic comparison of *R. torques* and *R. gnavus* based on PATRIC database analysis (67). *p≤0.05.
